# Genetic Monitoring and Molecular Phylogeny for Cymothoid Isopod Invasion in Egyptian Lake Qarun Fishes by Using adopted taxonomic marker (COI)

**DOI:** 10.1101/614578

**Authors:** Mohammed A. Hassan

## Abstract

Different types of aquatic ecosystem is abundant by a lot of crustaceans. There are species adapted to maximum of temperature, pressure and salinity. Isopods are a marine parasitic organism and commonly live in warmer seas. They are feeding on blood. Several species settle in the buccal cavity of fish. Parasite became a morbidity case in lake Qarun, GAFRD (General Authority for fish resources development, Fish Statistics yearbook, 2015). DNA barcoding gene (COI gene) was used to make the genetic characterization for the invasive species of Cymothoidae, *Cymothoidae sp* and recorded in the Genbank. Constructing a phylogeny depending on the degree of similarity between the new record (LC138010.1) and the other records of Cymothoidae species in the Genebank then the environmental conditions were compared according to the analysis of the constructed tree. Suez canal is the primary resource for the fingers of mullet which transferred to Qarun Lake (salinity, 49 ppt). Ballast water have an environmental effect by transferring the alien species in the Suez Gulf which have a warm weather. This study suggests that the Cymothoidae is expanded invasive species came from its habitat regions to a closed lakes which have a similar condition of water environment, temperature and depth to live and to be a morbidity in Lake Quran. Ballast water is a risky way to transfer the invasive species to Suez Canal then New Suez Canal poses and lead to biodiversity risks. Using **eDNA** techniques to diagnose the invasive species is very important unit which must be established in general authorities of fish resources and development in Egypt and other countries which invaded by different alien species.

## 1. Introduction

Lake Qarun (Birket Qarun), is known as an old natural lake in Egypt, located in Fayoum and the distance from Cairo is about 90 kilo meter southern Cairo. It is an inland closed sink of water with surface area of about 330 Km2, described by high salinity and the increasing of evaporation during summer [1]. The Cymothoidae is an ectoparasite which became a morbidity case in Lake Qarun in the summer of year 2015 [2]. The problem was gradually propagated reaching a terrible invasion causing fish mass loss, great marketing problem and drop of fish stock that lead to economic losses in addition to the impact on fishermen who working in this area [3]. The species of *Tilapia zillii* (important fishes are infested by Cymoithoids (Crustacean: Isopoda; Cymothoidae), Cymoithoids are hermaphrodites and nurturing on blood and different tissues of their suitable host [4]. Isopod species occupy the buccal cavity, inhabit the respiratory cavity or attached to the surface of body. The water salinity in Lake Qarun ranged between 42 and 49 ppt. and in Gulf of Suez ranged between 40 and 42 ppt. Water Temperature ranged between 25.5 and 35.7 °C in Lake Qarun but ranged between 29.3 and 33.4 °C in Gulf of Suez [5]. Shifting host species happened to Cymoithoids species and the molecular phylogeny for 27 Cymothoid species using 16s rDNA and COI gene recommended that Cymothoids may have originated in deep seas, then diffused to shallow seas, and then to brackish or freshwater [6]. Host specificity and host range vary among. In general, however, cymothoids that attach to external surfaces of fish display wider host ranges than other species, sometimes mount fish orders [6]. Several molecular phylogenetic studies on the family Cymothoidae, the reference frameworks are imprecise; vague sequences were included in previous phylogenies based on 16S rRNA and COI sequences [7]. Cymothoids that are parasitic in the buccal cavities show average distance accross host range [8]. The most of cymothiodae species occupy the Amazonia in South America. Several species of the genus *Ichthyoxenos* live in freshwater habitats of eastern and south-eastern Asia, and central Africa [9]. The invasion of Red Sea Biota into the Mediterranean by way of the Suez Canal [10]. In this study we focused in genetic monitoring method (DNA Barcoding Gene, COI gene) to characterize and classify a one species of Cymothoidae which invade the Lake Qarun during summer of 2016 that destroyed the fish stock of Qarun lake then a phylogeny was constructed depending on molecular basis evolution to measure the distance of homology by Maximum Likelihood algorithm and constructed the best and confidence phylogeny depending on bootstrapping algorithm. Transfer the fingers of *Mugil cephalus* from Suez Gulf [2]. The water environment in Qarun is similar to the environment of Suez Gulf; the water salinity in Lake Qarun ranged between 42 and 49 ppt. and in Gulf of Suez ranged between 40 and 42 ppt [5]. Water Temperature ranged between 25.5 and 35.7 C° in Lake Qarun but ranged between 29.3 and 33.4 C° in Gulf of Suez [5].

## 2. Material and Methods

### 2.1 Area of study and fish sampling

Lake Qarun is a natural saline lakes located in the lowest part of Fayoum Depression in the Western Desert of middle Egypt between the longitudes of 30° 24′ & 30° 49′ E and latitudes of 29° 24′ & 29° 33′ N [18]. A total of 23 fish samples belonging to *Tilapia zilli* were caught by fishermen from the lake Qarun in March, 2015. Isopods (the adult female cymothoid carried the developing embryos in the marsupium) that detected were isolated from gills arches and preserved in (0.5M *EDTA*, chelating agent) and Ethanol 70% and adjusted the pH of the buffer to be (pH 8.3). The isopods was washed by *18.2* MΩ·cm at 25 °C and stored in −80 C°. We collected one species of Cymothoidae sp. from the fishes, *Tilapia zilli*. The infested hosts were captured (Fig. 1), (Figs. 2 &3).

**Fig. 1.**
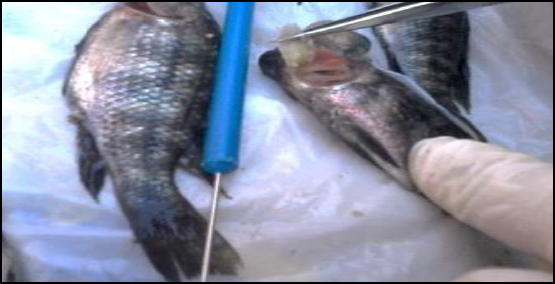
*Cymothoidae sp. collected from Tilapia* which located and attached to the gills.

**Fig. 2.**
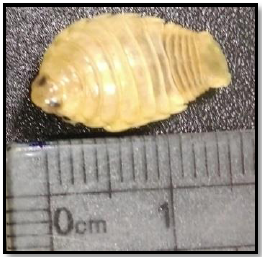
Dorsal view for *Cymothoidae sp.*, length = 1.5 cm.

**Fig. 3.**
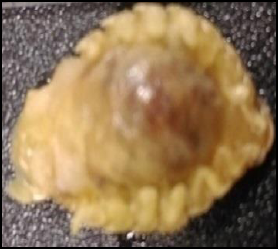
Ventral view for *Cymothoidae sp.*

### 2.2 DNA extraction

Two females of *Cymothoidae sp* homogenized by liquid nitrogen in 400 µl CTAB (hexadecyltrimethyl ammonium bromide) buffer (2% CTAB, 1.0 M NaCl, 75 mM EDTA (pH 8.0), 35 mM Tris-HCl (pH 8.0) containing 0.1% sodium dodecyl sulfate (SDS) and 0.2% beta-mercaptoethanol, followed by incubated at 65 °C for 1 h, according to the method of [15]. Proteinase K was added to the samples to a final concentration of 0.1 mg.m-1 and incubated the samples overnight at 37°C. then the DNA was extracted by using phenol-chloroform [15].

### 2.3 PCR and sequencing

PCR amplification for the the COI regions of mtDNA was performed by using the forward primer, LCO1490 (5′ ggtcaacaaatcataaagatattgg 3′) and reverse primer, HCO2198 (5′ taaacttcagggtgaccaaaaaatca 3′) according to [16]. PCR was performed in 50 μl reaction volume containing 25μl of Taq Master (2x), Jena bioscience, 1-2 μl each primer (0.05 µM for each primer) and DNA template (DNA concentration ≈35-50 ng/ μl). The thermal cycle profile was as follows: initial denaturation at 94 °C for 10 min; 30 cycles of 94 °C for 30 s, 48 °C for 1 min, and 72 °C for 1 min; and a final extension at 72 °C for 10 min. PCR products were purified from agarose gel using kit of jena bioscience (kiT# PP-202S) then subjected to direct cycle sequencing by using BigDye Terminator technology (Applied-Biosystems)

### 2.4 Sequence analysis and submission to *DDBJ*

The COI fragment of *Cymothoidae sp* which sequenced was submitted to DNA Data Bank of Japan under accession number (LC138010.1) and used as a query sequence to hit the database of gene bank then downloaded as local aligned sequences. All the steps of analysis started by a tool named open reading frame finder (ORFfinder), https://www.ncbi.nlm.nih.gov/orffinder/. Using translated table amino acid 5.0 of invertebrates which returned the range of each ORF (Open Reading Frames), along with its protein translation then the Blastp program was avaliable as an online server of Protein Blast (https://blast.ncbi.nlm.nih.gov/Blast.cgi) was used to find the homologues sequences which recorded in the Genbank. The Mafft sever for multiple sequences alignment (using the slow option of the algorithm, G-INS-1 (Slow; progressive method with an accurate guide tree). MEGA program ver 7.0 was used to Estimate Maximum-Likelihood Phylogenies: by using maximum likelihood ML statistical method and adjusted the value of bootstrapping option to be 500. The main steps were concluded in (Fig. 4).

**Fig. 4.**
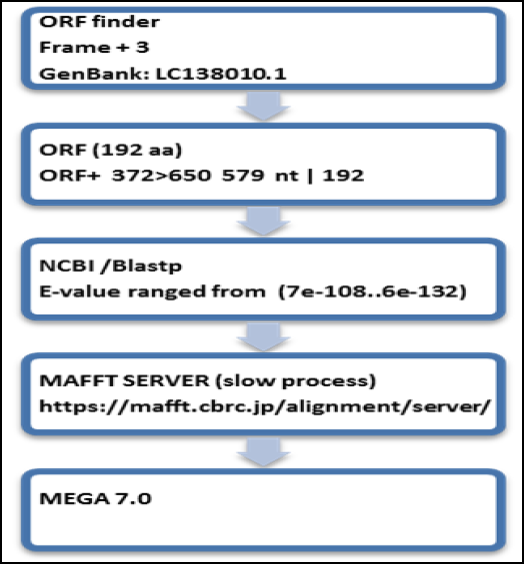
The main steps to construct the phylogeny.

### 2.5 sequence analysis to find pattern

The GenBank record (BAU78233.1), of protein sequence, cytochrome c oxidase subunit I, partial (mitochondrion) [Elthusa vulgaris] was a cross reference for the accession number (LC138010.1) nucleotide sequence and retrieved to hit the protein database sequence by using Blastp to find the homologues protein records(The Sequences which produced a significant alignments were 100 records of Cymothoidae isopods). The protein homologus sequences were aligned by MAFFT server to construct the multiple sequence alignment blocks where the longest and conserved block was copied and past from the MAFFT result of multiple sequence alignment by using Jalview viewer (version 2.10.5) [20] and exported to be as a fast format records of the detected conserved block. The exported fasta records of the conserved block were processed by an online server which used to expect a perect pattern of proteins fasta format records, Pratt algorithm of EBI analysis tool. The Pratt algorithm (https://www.ebi.ac.uk/Tools/pfa/pratt/) was used to find a specific pattern among the cymothiodae species.

## 3.0 Results and discussions

### 3.1 Expected size of PCR

The expected size of COI gene which produced by PCR was 672 and after trimming 20 bp (right and left sequences) of chromatogram curve the processed file of ABI file of the sequencer was 652 bp and then converted to fasta format file. The file was processed and annotated then submitted to gene bank under accession number (LC138010.1).

### 3.2 Construction of Cymoithoids phylogeny

In the present study, the tree was constructed depending on COI genes and the combined dataset. The translated frame of the accession number (LC138010.1) was used as a query sequence (192 a.a) to target the database of protein records of COI gene. The homologues amino acid sequences which gave significant score of pairwise alignment where the E-vales were ranged from (7e^−108^..6e^−132^) to indicate for high a similarity between the recorded sequences and the translated query sequence (LC138010.1). The local aligned amino acid sequences were downloaded and aligned by Mafft algorithm to produce MSA, Multiple Sequence Alignment. The output of multiple sequence alignment was used to construct the the tree by MEGA 7.0 as illustrated in (Fig. 5). The evolutionary history was inferred by using the Maximum Likelihood method based on the JTT matrix-based model. The bootstrap consensus tree inferred from 50 replicates. Was taken to represent the evolutionary history of the taxa analysed. Branches corresponding to partitions reproduced in less than 50% bootstrap replicates are collapsed. The percentage of replicate trees in which the associated taxa clustered together in the bootstrap test (50 replicates) are shown next to the branches. Initial tree(s) for the heuristic search were obtained automatically by applying Neighbor-Join and BioNJ algorithms to a matrix of pairwise distances estimated using a JTT model, and then selecting the topology with superior log likelihood value. The analysis involved 50 amino acid sequences. All positions containing gaps and missing data were eliminated. There were a total of 184 positions in the final dataset. Evolutionary. analyses were conducted in MEGA7 [14].

**Fig. 5.**
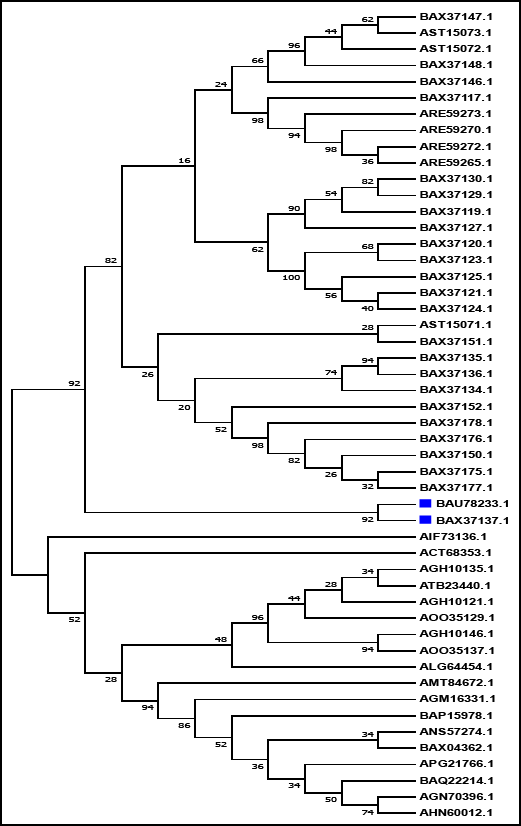
Molecular Phylogenetic analysis by Maximum Likelihood method.

### 3.3 phylogeny analysis and interpretation

The trees indicated that the invasive species (*Cymothoidae sp*) in Lake Qarun (BAU78233.1, translated in silicon of submitted accession number LC138010.1) was closely related to the Cymothoidae sp. Cy150091 which had the accession number (BAX37137). The environment in Lake Qarun was increased in salinity and reducing in dissolved oxygen where the DO was 3.5 ppm at depth 3m [4] and [5], which is a suitable environment for Elthusa sp. to be more in the lake. Marine cymothoids are almost limited occupant of shallow water, little of them being known from long depths [11]. The depth of the collected Cymothoidae from the shallower (eastern basin) Location of Qarun Lake was 7m [17] that was appoximately similar to the environments of (Cymothoidae sp. Cy150091, (BAX37137)). Cymothoidae sp. Cy150091 was located at 12 m in the location : unabe, Okinawa, JPN) [6]. The protein accession number was coded by its nucleotide accession number and annotated in the GenBank as follows : the nucleoitde accession number was translated by “/transl_table=5” and coded as follows, “/coded_by=LC138010.1: 45..>652”. All the previous annotations were used to annotate the accession number (LC138010.1) and produced the protein accession number (BAU78233.1). The depth of the shallower (eastern basin) Location of Qarun Lake, Fayoum, Egypt has a depth from 5m to 8.4m [17]. The main problem came from the Transferring the fingers of *Mugil cephalus* from Suez Gulf which infested by the Cymothoidae species. The water environment in Lake Qarun is similar to the environment of Suez Gulf; the water salinity in Lake Qarun ranged between 42 and 49 ppt. and in Gulf of Suez ranged between 40 and 42 ppt [5]. Water Temperature ranged between 25.5 and 35.7 C° in Lake Qarun but ranged between 29.3 and 33.4 C° in Gulf of Suez [5]. Ballast Water Bringing Invasive Species to Coasts. Maritime traffic could bring alien species as Cymothoidae alien species · New Suez Canal poses biodiversity risks for Malta [12]. Historically, the bio fouling of ships’ hulls has always been considered the oldest tools that causing the bringing of marine invasive species. in the last few years, studies have been concentrated mainly towards the introduction of non-indigenous species (invasive species) via ballast water [13].

### 3.4 Pattern of the highly conserved blocks

Overall, patterns of amino acid variation suggest convergent or parallel evolution at the protein level connected to the transition into a parasitic life style [19]. The aligned sequences (100 records of Cymothoidae which similar to the query sequence (BAU78233) produced by MAFTT sever where 100 sequences had 216 total sites, 198 gap-free sites and 197 conserved sites. The longest and highest conserved block had 23 amino acid resides and five different amino acids.

The consensus sequence in this block was as follows (LLLLSLPVLAGAITMLLTDRNLNTSFFDP) (Fig. 6).

**Fig. 6.**
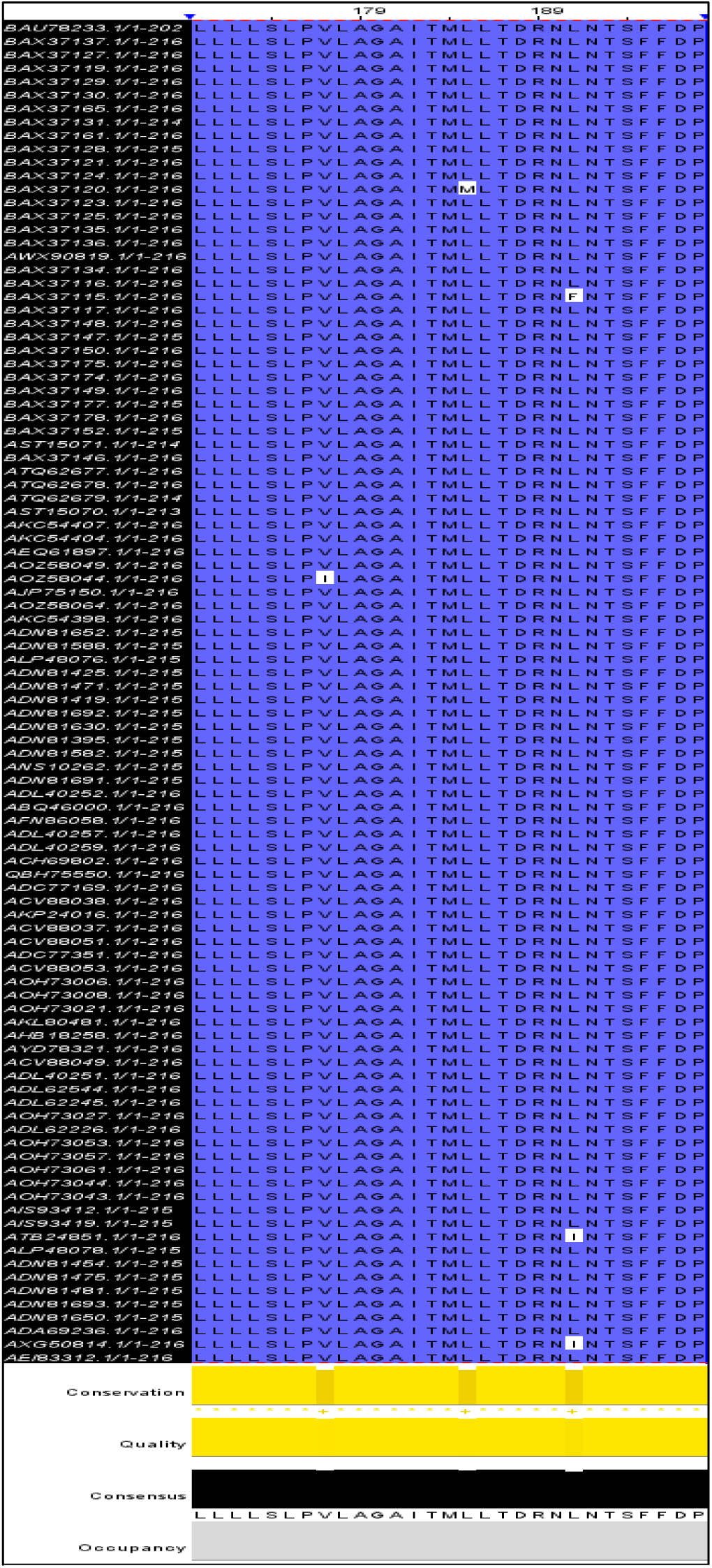
The conserved block between the cytochrome c oxidase subunit 1 which illustrated the different five amino acids across the conserved column.

The best pattern which produced by Pratt algorithm [21] was characterized and processed to be a best patterns With alignment (L-S-L-P-x-L-A-G-A-I-T-M-x-L-T-D-R-N-x-N-T-S-F-F-D-P).

## 4. Conclusion

Ballast Water Bringing Invasive Species to Coasts. Maritime traffic could bring alien species as Cymothoidae alien species · New Suez Canal poses biodiversity risks. Using eDNA techniques to diagnose the invasive species is very important unit which must be established in general authorities of fish resources and development in Egypt and other countries which invaded by different alien species. Searching for motifs which related to cytochrome c oxidase subunit 1 of Cymothoidae and linked to environment which invaded by marine isopods

## 5. Acknowledgement

My deepest thanks to the Chemist Mona Habib, the manager of central lab of General Authority For Fish Resources Development (GAFRD), Egypt and Dr., Doae Mohamed, Department of ecology and environmental research (GAFRD) to support me during the collection of the samples from Lake Qarun and supporting me by the environmental references.

## References

1. Gupta, G., & Abd El-Hamid, Z. (2003). Water quality of lake Qarun, Egypt. International journal of environmental studies, 60(6), 651–657.

2. General Authority for Fish Resources GAFRD, Cairo, Egypt. (2015). Fish Statistics year book.

3. Nisreen E Mahmoud, M M Fahmy and Mai Abuowarda, M (2017): An Investigation of Cymothoid Isopod Invasion in Lake Qarun Fishes with Preliminary Trial for Biological Control. International Journal of Chem Tech Research, Vol.10, No.12, pp 409–416.

4. Nisreen E. Mahmoud, M.M. Fahmy, Mai Abuowarda M and Marwa S. Khattab2 (2017): Parasitic Cymothoid Isopods and their Impacts in Commercially Important Fishes From Lake Qarun, Egypt. International Journal of Chem Tech Research, Vol.9, No.12, pp 221–229.

4. Gupta, G., & Abd El-Hamid, Z. (2003). Water quality of lake Qarun, Egypt. Intern. J. Environ. Studies, 60(6), 651–657.

5. Abdelaziz, M., Ibrahem, M. D., Ibrahim, M. A., Abu-Elala, N. M., & Abdel-moneam, D. A. (2017). Monitoring of different vibrio species affecting marine fishes in Lake Qarun and Gulf of Suez: Phenotypic and molecular characterization. The Egyptian Journal of Aquatic Research, 43(2), 141–146.

6. Hata, H., Sogabe, A., Tada, S., Nishimoto, R., Nakano, R., Kohya, N. & Kawanishi, R. (2017). Molecular phylogeny of obligate fish parasites of the family Cymothoidae (Isopoda, Crustacea): evolution of the attachment mode to host fish and the habitat shift from saline water to freshwater. Marine Biology, 164(5), 105.

7. Ketmaier V, Joyce DA, Horton T, Mariani S (2008) A molecular phylogenetic framework for the evolution of parasitic strategies in cymothoid isopods (Crustacea). J Zool Syst Evol Res 46:19–23.

8. Hadfield KA, Bruce NL, Smit NJ (2015) Review of Mothocya Costa in Hope 1851 Crustacea Isopoda Cymothoidae from southern Africa, with the description of a new species. Afr Zool 50:147–163.

9. Tsai, M. L., & Dai, C. F. (1999). Ichthyoxenus fushanensis, new species (Isopoda: Cymothoidae), parasite of the fresh-water fish Varicorhinus barbatulus from northern Taiwan. Journal of Crustacean Biology, 19(4), 917–923.

10. Por, F. D. “The Influx of Red Sea Biota into the Mediterranean by way of the Suez Canal, vol. 23.” (1978).

11. Brusca, R. C. (1981). A monograph on the Isopoda Cymothoidae (Crustacea) of the eastern Pacific. Zoological Journal of the Linnean Society, 73(2), 117–199.

12. Fuchs, C. (2008). Convention on International Trade in Endangered Species of Wild Fauna and Flora (CITES)-Conservation Efforts Undermine the Legality Principle. German LJ, 9, 1565.

13. Bouda, A., el Islam Bachari, N., Nacef, L., & Bensari, B. (2017). Risk Analysis of Invasive Species Introduction in the Port of Arzew, by Calculation of Biofouling Surface on Ships’ Hulls. Environmental Modeling & Assessment, 1–8. Chicago.

14. Sudhir Kumar, Glen Stecher, and Koichiro Tamura (2015) MEGA7: Molecular Evolutionary Genetics Analysis version 7.0. Molecular Biology and Evolution (submitted).

15. Ota, Y., Hoshino, O., Hirose, M., Tanaka, K., & Hirose, E. (2012). Third-stage larva shifts host fish from teleost to elasmobranch in the temporary parasitic isopod, Gnathia trimaculata (Crustacea; Gnathiidae). Marine biology, 159(10), 2333–2347.

16. Vrijenhoek, R. “DNA primers for amplification of mitochondrial cytochrome c oxidase subunit I from diverse metazoan invertebrates.” Mol Mar Biol Biotechnol 3.5 (1994): 294–9.

17. Abdel-Satar, A. M., & Sayed, M. F. (2010). Sequential fractionation of phosphorus in sediments of El-Fayum lakes—Egypt. Environmental monitoring and assessment, 169(1-4), 169–178.

18. Mohamed, F. A. (2009). Histopathological studies on Tilapia zillii and Solea vulgaris from Lake Qarun, Egypt. World Journal of Fish and Marine Sciences, 1(1), 29–39.

19. Pentinsaari, M., Salmela, H., Mutanen, M., & Roslin, T. (2016). Molecular evolution of a widely-adopted taxonomic marker (COI) across the animal tree of life. Scientific Reports, 6, 35275.

20. Waterhouse, A.M., Procter, J.B., Martin, D.M.A, Clamp, M. and Barton, G. J. (2009) Jalview Version 2 - a multiple sequence alignment editor and analysis workbench Bioinformatics doi: 10.1093/bioinformatics/btp033

21. Jonassen, I., Collins, J. F., & Higgins, D. G. (1995). Finding flexible patterns in unaligned protein sequences. Protein science, 4(8), 1587–1595.

